# Detection of *Mycobacterium tuberculosis* in pediatric stool samples using TruTip technology

**DOI:** 10.1101/564088

**Authors:** AW Mesman, M Soto, J Coit, R Calderon, J Aliaga, NR Pollock, M Mendoza, FM Mestanza, CJ Mendoza, MB Murray, L Lecca, R Holmberg, MF Franke

## Abstract

**Background:** Rapid and accurate diagnosis of childhood tuberculosis (TB) is challenging because children are often unable to produce the sputum sample required for conventional tests. Stool is an alternative sample type that is easy to collect from children, and studies investigating the use of stool for molecular detection of *Mycobacterium tuberculosis (Mtb*) have led to promising results. Our objective was to evaluate stool as an alternative specimen to sputum for *Mtb* detection in children. We did so using the TruTip workstation (Akonni Biosystems), a novel automated lysis and extraction platform.

**Methods:** We tested stool samples of 259 children who presented with TB symptoms, aged 0-14 years old, in Lima, Peru. Following extraction with TruTip, we detected the presence of *Mtb* DNA by IS6110 real-time PCR. We calculated assay sensitivity in two groups: (1) children with culture confirmed TB (N=22); and (2) children with unconfirmed, clinically diagnosed TB (N=84). We calculated specificity among children in whom TB was ruled out (N=153). Among children who were diagnosed with TB, we examined factors associated with a positive stool test.

**Results:** Assay sensitivity was 59% (95% confidence interval [CI]: 39%-80%) and 1.2% (95% CI: 0.0%-6.5%) in children with culture-confirmed and unconfirmed clinically-diagnosed TB, respectively, and specificity was 97% (95% CI: 93%-99%). The assay detected *Mtb* in stool of 7/7 children with smear-positive TB (100% sensitivity; 95% CI: 59%-100%), and in 6/15 of children with smear-negative, culture-confirmed TB (40% sensitivity; 95% CI: 16%-68%). Older age, smear positivity, culture positivity, ability to produce sputum and cavitary disease were associated with a positive stool result.

**Conclusion:** Testing of stool samples with the TruTip workstation and IS6110 amplification, yielded sensitivity and specificity estimates comparable to other tests such as Xpert. Future work should include detection of resistance using the TruTip closed amplification system and assay optimization to improve sensitivity in children with low bacillary loads..

## Background

The World Health Organization estimates that one million new pediatric tuberculosis (TB) cases and 194,000 childhood TB deaths occurred in 2017 (1). Rapid case detection and treatment initiation is critical to minimizing TB morbidity and mortality in children but is hampered by the absence of a rapid, accurate diagnostic tool for this group. Bacteriologic confirmation of *Mycobacterium tuberculosis (Mtb*) in children is often difficult to achieve because they are frequently unable to expectorate sputum for bacteriologic testing and often have paucibacillary disease that cannot be detected using sputum smear microscopy, culture, and/or molecular testing [e.g. Xpert (Cepheid, Sunnyvale CA, USA)]. Sputum induction and gastric aspiration can be used to obtain respiratory specimens from children unable to expectorate sputum; however, gastric aspiration is invasive and neither procedure is widely implemented in resource-constrained settings. Due to these diagnostic challenges, bacteriologic confirmation of TB is obtained in only a small minority of children diagnosed with TB (2–4).

Stool can be easily obtained from most children, and *Mtb* can be detected in stool using Xpert (5–10) or other laboratory-developed PCR assays (11–13). In the case of Xpert, the process is automated, but detection of drug resistance is currently limited to rifampin-associated resistance mutations in rpoB. Non-integrated merthods for DNA, isolation and amplification of DNA using extraction kits and in-house tests are often laborious, multistep procedures. An ideal test would be an automated point-of-care workstation with integrated capacity for both *Mtb* and expanded drug resistance testing-criteria listed in the target product profile for novel TB diagnostics in low resource settings (14). The TruTip workstation is an automated platform including lysis and homogenization with TruTip nucleic acid extraction and purification (Akonni Biosystems, Frederick, MD, USA) (15–17). TruTip has been used for nucleic acid isolation from a variety of pathogens and sample types and has demonstrated efficient *Mtb* DNA recovery from raw sputum (15,18). The platform can be connected to a closed amplicon system for amplification and microarray-based detection of *Mtb* as well as a number of drug resistance-associated mutations (17,19–21). The aim of the present study was to estimate sensitivity and specificity of *Mtb* detection in stool from children with symptoms compatible with intrathoracic TB in Lima, Peru, using this novel technology with IS6110 real-time PCR.

## Methods

### Ethics

Study participants’ guardians provided written informed consent to participate, and children eight years of age and older provided written assent. Consent for publication was not applicable. All study procedures were approved by the Ethics Committee of Peru’s National Institute of Health and the Office of Human Research Administration at the Harvard Medical School.

### Study population

Between May 2015 and February 2018, we recruited children to participate in a pediatric TB diagnostics study. Eligible children were less than 15 years of age, had a history of contact with an adult with TB within the previous two years, and presented to a participating public sector health center in Lima, Peru with symptoms compatible with TB (i.e., persistent cough for more than two weeks; unexplained weight loss; unexplained fever for more than one week; and/or unexplained fatigue or lethargy) (22). For this analysis, we included the subset of children with culture-confirmed TB or clinically-diagnosed unconfirmed TB who had at least one stool sample available. We also included a subset of the children in whom TB had been ruled out (i.e, controls), matching on age and sample collection date when possible.

### Study procedures and sample collection

Children were evaluated for TB per Peruvian National Tuberculosis Strategy guidelines (23). In brief, children provided up to two gastric aspirate (GA) and/or sputum samples (expectorated or induced) for smear and culture, and Ministry of Health pediatric pulmonologists considered these results as well as medical history, physical examination, chest X-ray findings and tuberculin skin testing (TST) results to diagnose or rule out TB. GA samples were neutralized to a pH of 6.8-7.2 upon collection. We requested two stool samples from all children for research purposes. From children who were diagnosed with TB, we aimed to collect these samples prior to TB treatment initiation. Stool collection took place at home or the health center. For children in diapers, plastic wrap was inserted on the inside of the diaper and a urine collection bag was attached to the child. The latter served to collect a urine sample as well as to prevent mixing of urine and stool.

Following collection, all samples were transported by cold-chain to the laboratory, aliquoted and stored at −80°C until testing. Because testing of stool did not occur in real time and because the assay was experimental, stool testing results were not shared with families or healthcare providers. HIV testing was not performed because the HIV prevalence among children in the study setting is <0.1% (24).

### Laboratory procedures

Sample processing and testing was performed in the BSL-3 and BSL-2 facilities at the Socios En Salud Sucursal Laboratory in Lima. We centrifuged and decontaminated sputum and GA samples in 2% NaOH/0.25% n-acetyl-L-cysteine (NALC). GA and sputum samples were analyzed by Ziehl-Neelsen staining/smear microscopy and liquid culture (BACTEC MGIT 960, BD Franklin Lakes, USA).

For lysis and DNA extraction procedures using the TruTip workstation we followed methods similar to previously described for sputum samples (18). In brief, 500 milligrams of thawed stool was homogenized in TE buffer, in a magnetically-induced vortexing (MagVor) element, and after a 20 minute heating step at 56°C, the samples were transferred to the TruTip workstation for DNA extraction (16). *M*t*b* DNA was amplified in duplicate by real-time PCR using a Roche 480 Lightcycler (Basel, Switzerland) with primers targeting the insertion element IS6110 (25). We considered a sample positive for *Mtb* DNA if, for both PCR replicates, the cycle threshold (Ct) value was <37 and the change in fluorescence was >6 units. We repeated the PCR step in duplicate for discordant replicates and considered the sample to be indeterminate for *Mtb* DNA if the two replicates remained discrepant after a second PCR run. Laboratory staff were blinded to participants’ clinical status.

### Data analysis

Data were analyzed using SAS (Version 9.4. SAS Institute Inc., Cary, NC, USA). We calculated sensitivity and specificity and 95% confidence intervals (CI) both at the child-level (including all samples from each child) and at the sample-level. Sensitivity of *Mtb* detection from stool was calculated separately for children with confirmed TB (i.e., based on a positive culture from sputum or GA) and clinically-diagnosed unconfirmed TB. Specificity was estimated among those in whom TB had been ruled out. In per-sample analyses we adjusted confidence intervals for clustering from multiple samples per child. Among all children with a TB diagnosis, we used chi-square tests to examine the associations between a positive stool test and the following variables: age, sex, smear and culture results, ability to spontaneously expectorate sputum, and cavitation on chest x-ray. Numerical precision for all percentages, p-values and 95% CIs is presented based on the evidence-based recommendations proposed by Cole (26).

## Results

### Study population

From 628 children enrolled in the study, we selected 259 children based on stool sample availability (22 children with culture confirmed TB, 84 children with unconfirmed, clinically diagnosed TB, and 153 children in whom TB had been ruled out (Fig 1)). These children had a median age of 5.1 years and 48% were female. Nearly all children (255/259; 98%) had at least one respiratory sample collected. Of the four with no respiratory sample, three were diagnosed based on clinical criteria and TB was ruled out in one. Children between the ages of 11 and 14 years old were more likely than younger children to have a culture-confirmed diagnosis, and all seven children with smear-positive sputum belonged to this older age group (Table 1). Thirteen of 22 children with confirmed TB were diagnosed based on expectorated sputum analysis and the remaining nine on gastric aspirate analysis.

**Table 1:** Demographic characteristics of children, stratified by TB status.

| Variable |  | All children<br>(N=259)<br>n (%) | SM+, CU+<br>TB<sup>a</sup><br>(N=7)<br>n (%) | SM-, CU+<br>TB<sup>a</sup><br>(N=15)<br>n (%) | All CU+ TB<br>(N=22)<br>n (%) | Unconfirmed<br>clinically<br>diagnosed TB<br>(N=84)<br>n (%) | No TB<br>diagnosis<br>(N=153)<br>n (%) |
| --- | --- | --- | --- | --- | --- | --- | --- |
| Age group<br>(years) | 0-4 | 125 (48) | 0 (0) | 8 (53) | 8 (36) | 42 (50) | 75 (49) |
|  | 5-10 | 37 (14) | 0 (0) | 5 (33) | 5 (23) | 34 (40) | 58 (38) |
|  | 11-14 | 97 (37) | 7 (100) | 2 (13) | 9 (41) | 8 (10) | 20 (13) |
| Female<br>sex |  | 124 (48) | 5 (71) | 6 (40) | 11 (50) | 47 (56) | 66 (43) |

<sup>a</sup>CU: culture, SM: smear

**Figure 1.**
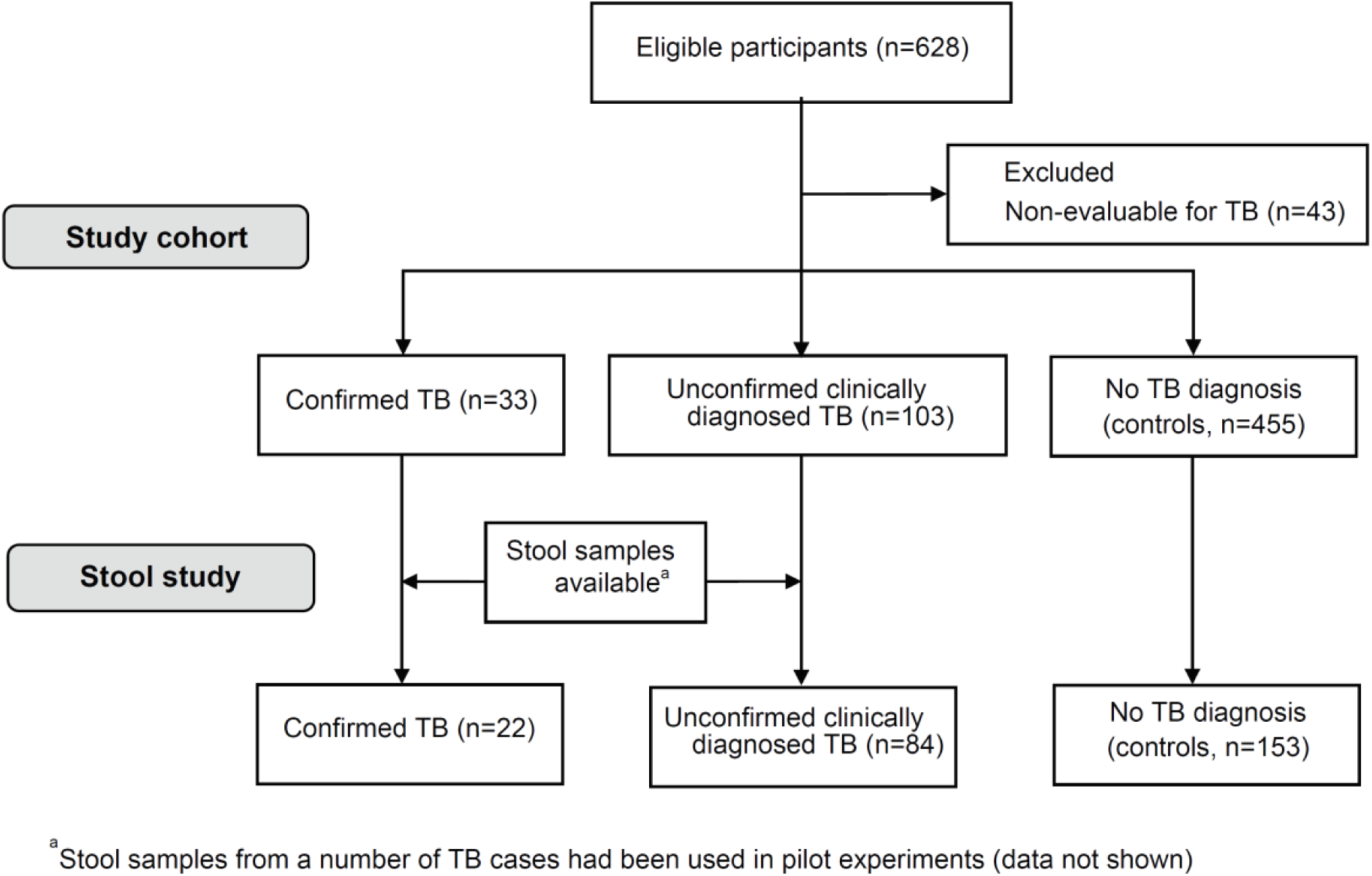
Flowchart of cohort enrollment and stool study inclusion.

From the 259 children, we collected 484 stool samples. Most children had two stool samples; however 34 children (13%) had only one sample available. Seventeen samples from children diagnosed with TB were collected after TB treatment initiation (median time from treatment initiation to collection: 3 days; range [1 – 11 days]). Eight samples were collected 72 hours or more after treatment start. Four of these eight were in children with smear-positive sputum because they were eligible to initiate TB treatment immediately, resulting in a narrow window between enrollment and treatment initiation during which pre-treatment samples could be collected.

### Per sample analyses

The assay detected *Mtb* in 25/484 samples. Only one sample, from a child in the control group, had an indeterminate result, which we excluded from further analyses. Among 13 stool samples from seven children with smear positive sputum, 100% (95% confidence interval [CI]: 59%-100%) were positive by the stool assay (Table 2). Sensitivity was lower in the smear-negative culture-confirmed group (23%; 95% CI: 5.6%-41%, Table 2). We detected *Mtb* in 4/282 (1.4%) samples from control children, resulting in a specificity of 99% (95% CI: 97-99.8%).

**Table 2:** Per sample analyses of *Mtb* stool assay in 259 children.

|  | Positive stool result (n/N) | Indeterminate stool result (n/N) | Sensitivity (95% CI<sup>a</sup>) | Specificity (95% CI<sup>a</sup>) |
| --- | --- | --- | --- | --- |
| All samples | 25/484 | 1/484 |  |  |
| Confirmed TB (SM+)<sup>b</sup> | 13/13 | 0/13 | 100% (59%-100%) |  |
| Confirmed TB (SM-) | 7/30 | 0/30 | 23.3% (5.6%-41%) |  |
| Confirmed TB (all smear grades) | 20/43 | 0/43 | 47% (27%-66%) |  |
| Unconfirmed, clinically diagnosed TB | 1/159 | 0/159 | 0.6% (0.00%-1.9%) |  |
| No TB diagnosis | 4/282 | 1/282 |  | 99% (97%-100%) |

<sup>a</sup>CI: confidence interval; 95% CI are adjusted for multiple samples per patient. <sup>b</sup>SM: smear

### Per child analyses

In child-level analyses, sensitivity of the stool assay conducted in each child’s first sample was 55% (95% CI: 33%-75%). When we included the results of both samples, we identified one additional confirmed TB case, which increased sensitivity to 59% (95% CI: 39%-80%) (Table 3). While the majority of children with a positive stool sample also had a spontaneous sputum sample that was culture positive (11/14, 79%), we detected *Mtb* in the stool of two children in whom TB was confirmed based on gastric aspirate and one child with unconfirmed, clinically diagnosed TB (this child had one expectorated and one induced sputum sample, both of which tested negative for *Mtb*). All three children were under five years of age.

**Table 3:** Child-level analyses of *Mtb* stool assay in 259 children.

|  |  | Positive stool result (n/N) | Sensitivity (95% CI<sup>a</sup>) | Specificity (95% CI) |
| --- | --- | --- | --- | --- |
| First sample | All children | 16/259 |  |  |
|  | Confirmed TB (SM+)<sup>b</sup> | 7/7 | 100% (59%-100%) |  |
|  | Confirmed TB (SM-) | 5/15 | 33% (13%-61%) |  |
|  | Confirmed TB (all smear grades) | 12/22 | 55% (33%-75%) |  |
|  | Unconfirmed, clinically diagnosed TB | 1/84 | 1.2% (0.0%-6.5%) |  |
|  | No TB diagnosis | 3/153 |  | 98% (94%-99.5%) |
| Both samples | All children | 18/259 |  |  |
|  | Confirmed TB (SM+) | 7/7 | 100% (59%-100%) |  |
|  | Confirmed TB (SM-) | 6/15 | 40% (16%-68%) |  |
|  | Confirmed TB (all smear grades) | 13/22 | 59% (36%-79%) |  |
|  | Unconfirmed, clinically diagnosed TB | 1/84 | 1.2% (0.0%-6.5%) |  |
|  | No TB diagnosis | 4/153 |  | 97% (93-99.3%) |

<sup>a</sup>CI: confidence interval, <sup>b</sup>SM: smear

### Factors associated with stool test positivity among TB cases

Features of adult-like TB (i.e., a positive smear result, a positive culture result, the ability to spontaneously expectorate sputum and cavitary TB), were associated with detection in stool and more common in the 11-14 year age group (Table 4). There was no association between stool positivity and sex.

**Table 4:** Factors associated with *Mtb* stool assay positivity among children with culture-confirmed or unconfirmed, clinically diagnosed TB (N=106).

| Variable |  | Positive stool result (N=14)<br>n (%) | Negative stool result (N=92)<br>n (%) | P-value |
| --- | --- | --- | --- | --- |
| Age group (years) | 0-4 | 3 (21) | 47 (51) | 0.0008<sup>a</sup> |
|  | 5-10 | 4 (29) | 35 (38) |  |
|  | 11-14 | 7 (50) | 10 (11) |  |
| Female sex |  | 7 (50) | 51 (55) | 0.8<sup>b</sup> |
| Positive smear result |  | 7 (50) | 0 (0.0) | <0.0001<sup>b</sup> |
| Positive culture result |  | 13 (93) | 9 (9.8) | <0.0001<sup>b</sup> |
| Cavitary disease (x-ray) |  | 5 (36) | 0 (0.0) | <0.0001<sup>b</sup> |
| Ability to produce sputum |  | 12 (86) | 33 (36) | 0.0006<sup>a</sup> |

<sup>a</sup>Chi-square test <sup>b</sup>Fisher's Exact test

## Discussion

We found that our stool test was 100% sensitive for *Mtb* detection in stool among children with smear-positive confirmed TB and 40% sensitive in children with smear-negative confirmed TB. Importantly, we detected *Mtb* in stool of three children below the age of five that were not able to expectorate sputum (two of whom were diagnosed based on GA samples and one who was culture negative by sputum, but clinically diagnosed). The sensitivity (59%) and specificity (97%) of this assay relative to liquid culture of respiratory samples was comparable to previous studies using Xpert for stool testing in children (7,27). Unfortunately, similar to other studies (6,28–30), we rarely detected TB in children with culture negative, clinically diagnosed TB.

Overall there is room for optimization to improve the sensitivity of smear-negative TB cases. Considerations for assay optimization include testing of multiple sample aliquots, incorporating an initial homogenization step and altering the sample volume. Our stool test protocol required a relatively small input volume (500 mg), which may minimize the impact of PCR inhibitors in stool that can impede the detection of *Mtb*. In fact, several studies show reduced sensitivity with larger sample volumes. For example, Banada et al reported higher sensitivity and fewer invalid test results when stool input volume was decreased from 1.2 to 0.6 grams (5). Recently, DiNardo et al obtained 67% sensitivity with an in-house developed using only 50 mg, demonstrating that even smaller volumes can be sufficient to reach high sensitivity.

In line with previous reports (8,31) we found strong associations between a positive stool test and features of more adult-like TB (i.e., cavitary disease; positive sputum culture; positive sputum smear; the ability to produce sputum), which were most common in the oldest age group. This emphasizes the impact of age and smear status on the outcomes of stool testing and has critical implications for the reporting of pediatric TB diagnostic studies because stool assay sensitivities may vary widely based on the percentage of children in the cohort with adult-like TB and/or high bacterial burden. Because children with adult-like TB are often able to produce a sputum sample for testing, a stool assay will only begin to fill the pediatric TB diagnostic gap if it is sensitive in children who cannot already spontaneously expectorate sputum. To facilitate an understanding of stool assay performance and its utility across settings, we recommend stratifying stool sensitivity results by some measure of disease severity, ideally smear status, and reporting the added yield among children unable to spontaneously expectorate sputum.

The TruTip workstation had a number of attractive attributes for *Mtb* DNA extraction from stool. Extraction procedures required limited pipetting steps compared to commercially available kits. Additional future advantages include that the extracted DNA eluate can be isolated for other purposes, such as sequencing. Most importantly, the TruTip workstation can be integrated with a microarray-based detection of *Mtb* as well as a number of drug resistance-associated mutations (17,19,20), which is a focus of future studies.

A limitation of our study was that we lacked follow-up data on the clinical evolution of children, a key criterion for classifying pediatric cases of clinically diagnosed, unconfirmed TB. To the extent that unconfirmed clinical TB was overdiagnosed, sensitivity will be underestimated in this group. Similarly, missed TB diagnoses among children in whom TB was ruled out could lead to an underestimate of assay specificity. The influence of these potential biases appears limited given the very low sensitivity among children with unconfirmed clinical TB and a high overall specificity. A second limitation of our study is the absence of children living with HIV. Studies have shown that sensitivity of stool assays may be higher in children living with HIV (6,7), in particular with severe immunosuppression (27), therefore, our results may not be generalizable to children living with HIV.

## Conclusion

In conclusion, the results of our study show that we can detect *Mtb* in pediatric stool samples. For detection we used the TruTip workstation in combination with realtime IS6110 PCR, and its performance is comparable to other platforms. Future work should include detection of resistance in stool samples using the TruTip closed amplification system and assay optimization to improve sensitivity in children with low bacillary loads.

## Abbreviations

AFB: acid fast bacilli
Ct: cycle threshold
DST: drug sensitivity testing
Mtb: Mycobacterium tuberculosis
PCR: Polymerase Chain Reaction
TB: tuberculosis
TST: tuberculin skin test
WHO: World Health Organization

## Declarations

### Ethics Approval and Consent to participate

Study participant guardians’ provided written informed consent, and children eight years of age and older provided written assent. All study procedures were approved by the Ethics Committee of Peru’s National Institute of Health and the Office of Human Research Administration at the Harvard Medical School.

### Consent for publication

Our manuscript does not contain individual person’s data. Consent for publication was not applicable.

### Availability of data and material

Reported data are available on request to the corresponding author.

### Author’s contributions

AM: led the data analysis and interpretation and wrote the manuscript. MS: implementation of the study and interpretation of results. JC: analysis and interpretation of results. JA: implementation and data analysis. RC: study design and data analysis. NP: study design and interpretation of results. MM: implementation of the study. FM: implementation of the study. CM: implementation of the study. MM: study design and interpretation of results, LL: study design and interpretation of results. RH: study design, implementation and interpretation of results. MF conceptualized and designed the study, interpreted the results, and contributed to writing and editing. All authors critically reviewed the manuscript and approved the final version.

## Acknowledgements

We are grateful for all children and guardians who participated in this study, and for the participating health centers of the Ministry of Health of Peru. We thank Dr. Rafael Alejandro Ortiz Sousa, Dr. Javier Nicolaz Jugo Rebaza, Dr. Ildauro Aguirre Sosa, Dra. Silvanna De la Gala De los Santos, Dr. Hernán Del Castillo Barrientos, Cynthia Pinedo Chuquizuta, Helen Marín Samanez and Gissela Valderrama Yalán for their participation and Zibiao Zhang for data management support. We thank Socios En Salud for contribution and dedication to health equity and all the health care workers who contribute to this study and treating patients.

